# Proteome encoded determinants of protein sorting into extracellular vesicles

**DOI:** 10.1101/2023.02.01.526570

**Authors:** Katharina Waury, Dea Gogishvili, Rienk Nieuwland, Madhurima Chatterjee, Charlotte E. Teunissen, Sanne Abeln

**Affiliations:** Department of Computer Science, Vrije Universiteit Amsterdam, Amsterdam, The Netherlands; Laboratory of Experimental Clinical Chemistry, Department of Clinical Chemistry, Amsterdam UMC, Location AMC, University of Amsterdam, Amsterdam, The Netherlands; Vesicle Observation Centre, Amsterdam UMC, Location AMC, University of Amsterdam, Amsterdam, The Netherlands; German Center for Neurodegenerative Diseases (DZNE), Bonn, Germany; Neurochemistry Laboratory, Department of Clinical Chemistry, Amsterdam Neuroscience, Amsterdam UMC, Location VUmc, Vrije Universiteit Amsterdam, Amsterdam, The Netherlands; Centrum Wiskunde & Informatica, Amsterdam, The Netherlands

## Abstract

Extracellular vesicles (EVs) are membranous structures released by cells into the extracellular space and are thought to be involved in cell-to-cell communication. While EVs and their cargo are promising biomarker candidates, protein sorting mechanisms of proteins to EVs remain unclear. In this study, we ask if it is possible to determine EV association based on the protein sequence. Additionally, we ask what the most important determinants are for EV association. We answer these questions with explainable AI models, using human proteome data from EV databases to train and validate the model. It is essential to correct the datasets for contaminants introduced by coarse EV isolation workflows and for experimental bias caused by mass spectrometry. In this study, we show that it is indeed possible to predict EV association from the protein sequence: a simple sequence-based model for predicting EV proteins achieved an area under the curve of 0.77±0.01, which increased further to 0.84±0.00 when incorporating curated post-translational modification (PTM) annotations. Feature analysis shows that EV associated proteins are stable, polar, and structured with low isoelectric point compared to non-EV proteins. PTM annotations emerged as the most important features for correct classification; specifically palmitoylation is one of the most prevalent EV sorting mechanisms for unique proteins. Palmitoylation and nitrosylation sites are especially prevalent in EV proteins that are determined by very strict isolation protocols, indicating they could potentially serve as quality control criteria for future studies. This computational study offers an effective sequence-based predictor of EV associated proteins with extensive characterisation of the human EV proteome that can explain for individual proteins which factors contribute to their EV association.

## Introduction

Extracellular vesicles (EVs) are a heterogeneous group of lipid-delimited vesicles that are released by cells into the extracellular space [1, 2]. EVs have been observed across a manifold of cell types and all domains of life highlighting their universal biological importance [3]. On the basis of their function, cargo, size, and excretion pathways, EVs may be divided into three main types: exosomes, microvesicles, and apoptotic bodies [2].

Biomolecules associated with EVs are strong biomarker candidates as they can be isolated from easily accessible body fluids and provide insight into the state of the donor cells [4]. The clinical potential of EV biomarkers has been elucidated for several pathologies [5–7], especially for neurodegenerative diseases [4, 8]. Thus, there is a strong need to better understand the underlying mechanisms of protein association with EVs.

However, the investigation of EVs is hampered by the difficulties of correctly isolating and concentrating EVs out of complex body fluids. Although experimental guidelines are being established [9–11], there is a continued discussion about the optimal workflow, and many published studies do not meet the minimal criteria of EV enrichment and isolation [10]. Thus, the standardization of isolation and identification techniques is still a major hurdle, despite its importance for successful EV biomarker development [12, 13]. EV studies are prone to include non-EV contaminants during sample collection, isolation, concentration, and characterisation of EVs [13]. For instance, isolation kits, such as ExoQuick, might enrich EVs, but results in various contaminants, including antibodies and polymers [10]. It has been suggested that more than 70% of all particles isolated from blood plasma are not EVs and the major contaminants are lipoprotein particles [14, 15] along with platelets [16]. For studies investigating the presence and absence of unique proteins in EVs, it is of particular importance that the number of unique contaminants in the data is as low as possible.

Continuous advancements in research have substantially contributed to EV proteome characterization. Vesiclepedia [17] and ExoCarta [18,19] are two major manually curated databases compiling identified EV cargo. While these two databases provide a rich information source, they must be treated with caution and analysed carefully considering major types of artifacts likely to be present in the data.

It is crucial to be aware of how reported proteins may be associated with EVs. Proteins listed in these databases can be (i) inside the vesicles (”cargo”), (ii) on the outside (”protein corona”), as well as (iii) bound to the membrane. Thus, we use the term *EV association* and collectively refer to such proteins as EV proteins.

For biomarker research, all types of EV association may be relevant as the presence of a biomarker candidate within the EV lumen or membrane might explain its limited or missing detection, e.g., using immunoassays [20]. Knowledge of EV association can provide directions for experimental analysis, *i.e*., the need for EV isolation and disruption techniques prior to protein detection. Additionally, the systematic analysis of EV proteins might provide valuable insights into their characteristics and thereby can increase our understanding of the cell’s sorting mechanisms and of the function of EVs.

While there have been numerous attempts to predict subcellular [21], as well as extracellular matrix [22,23] localization of proteins based on machine learning approaches, prediction of EV localization has been pursued much less. Ras-Carmona *et al*. attempted the prediction of protein secretion by EVs but limited their study to exosome cargo proteins. They reported an area under the curve (AUC) of 0.76±0.03 using dipeptide composition features [24]. In this study, we focus on EV association predictions for the human proteome. In addition, we put an emphasis on the explainability of the machine learning models, both to reveal general sorting trends, as well as to identify potential mechanisms for individual proteins.

There are two primary aims of this study: 1. To ascertain the possibility to predict EV association based on amino acid sequence using machine learning; 2. to investigate if the presence or absence of a specific human protein in EVs is associated with the sequence, physicochemical, and structural features of this protein, as well as post-translational modification (PTM) annotations.

We analyzed publicly available human EV protein data and corrected these for a potential detection bias by excluding proteins not recognized in mass spectrometry (MS) studies and proteins identified by unreliable EV isolation workflows. We constructed a wide variety of informative protein properties as input for the machine learning models; all these properties could either be calculated directly from the protein sequence or were derived from curated database annotations. These handcrafted features allowed us to interpret the machine learning model predictions, and link them to potential EV sorting mechanisms. This study offers an effective sequence-based predictor of “a ticket to a bubble ride” [25] and an extensive and systematic characterisation of the human EV proteome.

## Results

In this study, we aimed to answer if prediction of EV association is a feasible task. To obtain annotations suitable to train such a predictor, we designed a data curation workflow combining several resources and filtering steps as seen in Figure 1 and described in detail in *Methods*. Subsequently, we trained a machine learning model and identified the properties most characteristic of EV and non-EV proteins both at a global level, and for individual proteins.

**Figure 1.**
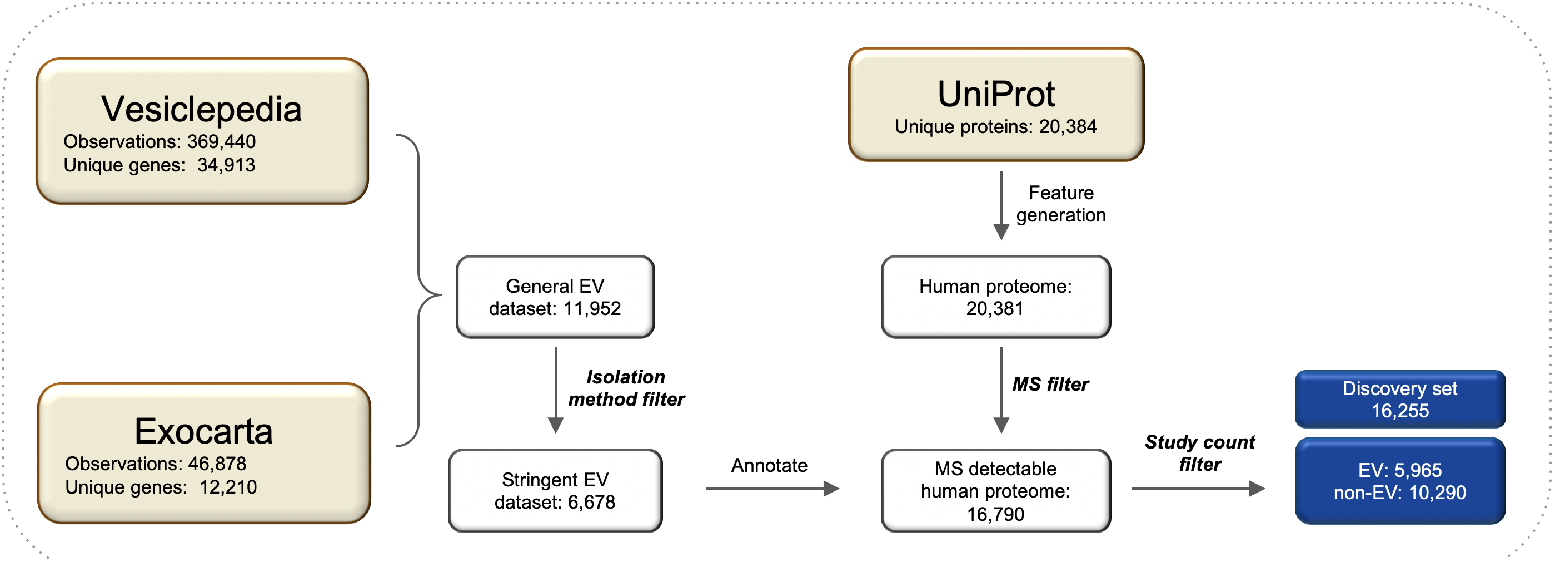
Data curation workflow. Squares represent datasets with remaining entries. Three datasets from databases Vesiclepedia, ExoCarta and UniProt are colored in khaki. Unique proteins from Vesiclepedia and ExoCarta were merged to construct a General EV dataset, and proteins identified by unreliable isolation workflows were removed to obtain the Stringent EV dataset. Sequence-based features as well as annotations were generated for each protein in the human proteome. Human proteins not detectable by MS were removed by the MS filter. All unique MS-detectable human proteins were annotated regarding their EV association using the stringent EV dataset. Lastly, rarely detected EV proteins (count ≤ 2) were removed from the dataset entirely resulting in the EV-annotated discovery set (blue). EV - extracellular vesicle, MS - mass spectrometry.

### EV proteomics data requires extensive data filtering steps

Several filtering steps were incorporated into the EV data curation workflow to reduce the likelihood of contaminants and any systematic bias introduced by mass spectrometry (MS); this workflow led to the EV-annotated discovery dataset (Figure 1). Additionally, we examined EV-annotated datasets that skip some of the filtering steps to determine what properties are truly connected to EV sorting processes and which might be a manifestation of biases; the data curation for these datasets is shown in Figure S1.

While MS is most suitable for the broad detection of proteins, it creates experimental data biased towards proteins of higher molecular weight and lower isoelectric point [26]. As most EV database entries have been identified by MS, we wished to investigate and correct for bias introduced by MS. We analysed the effect of the *MS filter*, as described in *Methods*, by comparing molecular weight distributions in the datasets with and without this filtering step. MS clearly struggles to detect low molecular weight proteins within the human proteome as especially low molecular weight proteins are missing in the MS-detectable proteome compared to the full human proteome (Figure 2A). This confirms previous results [26]. This bias is strongly pronounced in the unfiltered EV-annotated dataset (Figure 2B). Here, the non-MS-detectable subset of the human proteome is annotated as non-EV which explains the low molecular weight peaks in the non-EV proteome. When applying the MS filter and, thus, using only the MS-detectable subset of the human proteome, the discrepancy in molecular weight between EV and non-EV associated proteins is much smaller (Figure 2C), illustrating that the constructed filter successfully corrects for technical bias introduced by MS. While we only show the effects of the filtering step in terms of protein size, we assume this filter will also help to correct any additional unknown MS detection biases.

**Figure 2.**
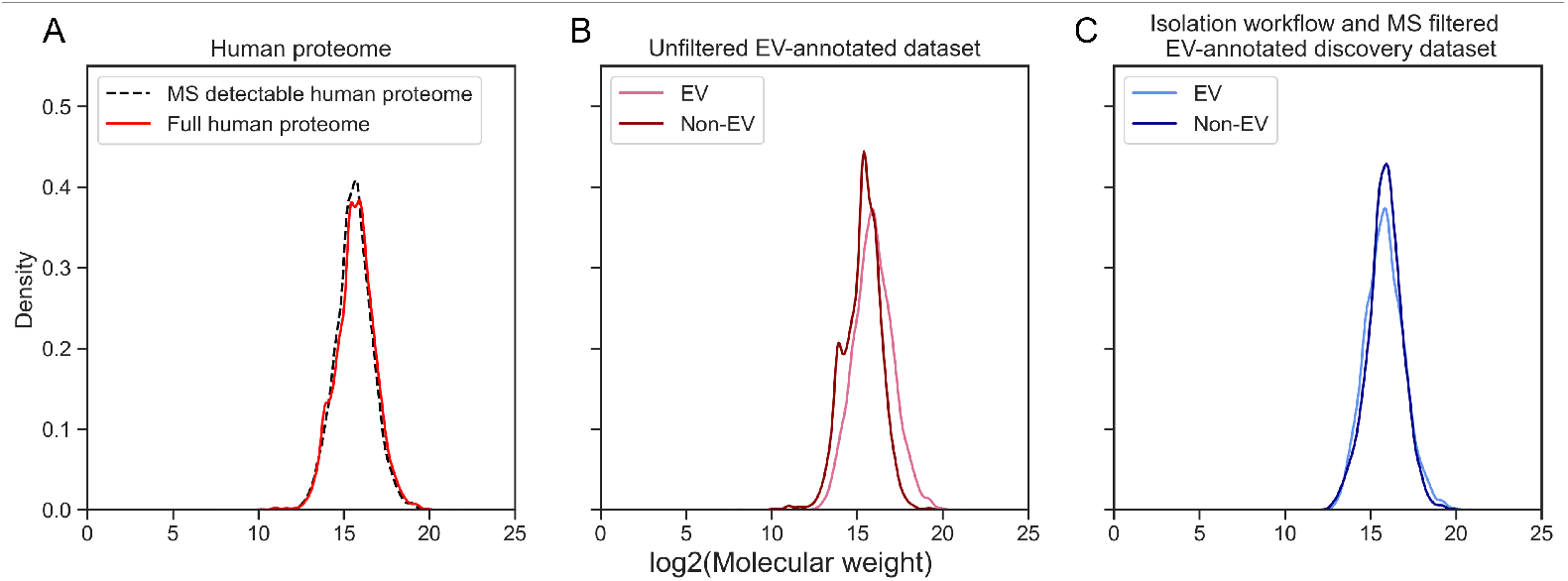
Density plots of log2-transformed molecular weight across the human proteome and EV-annotated datasets. (**A**) Distribution of log2-transformed molecular weight of the MS-detectable human proteome compared to the full human proteome. MS struggles to detect low molecular weight proteins of the human proteome. (**B**) The molecular weight densities of EV and non-EV proteins in the unfiltered EV-annotated dataset highlight the discrepancy in molecular weight between EV and non-EV proteomes. (**C**) The much more similar molecular weight distribution of EV and non-EV group show how the MS filter step diminishes the experimental bias introduced by MS. EV - extracellular vesicle, MS - mass spectrometry.

### Prediction of EV associated proteins is feasible

To investigate if there is a signal in a protein’s sequence that determines its EV association, we constructed machine learning models to predict this property. The sequence properties of a protein are the input features for the model; the output is a prediction if the protein is EV associated or not. Two different random forest (RF) models were trained on the discovery dataset: (i) incorporating only sequence-based features; (ii) incorporating sequence-based features and curated annotations. Figure 3A displays their respective receiver operating characteristic (ROC) curves and AUC scores.

**Figure 3.**
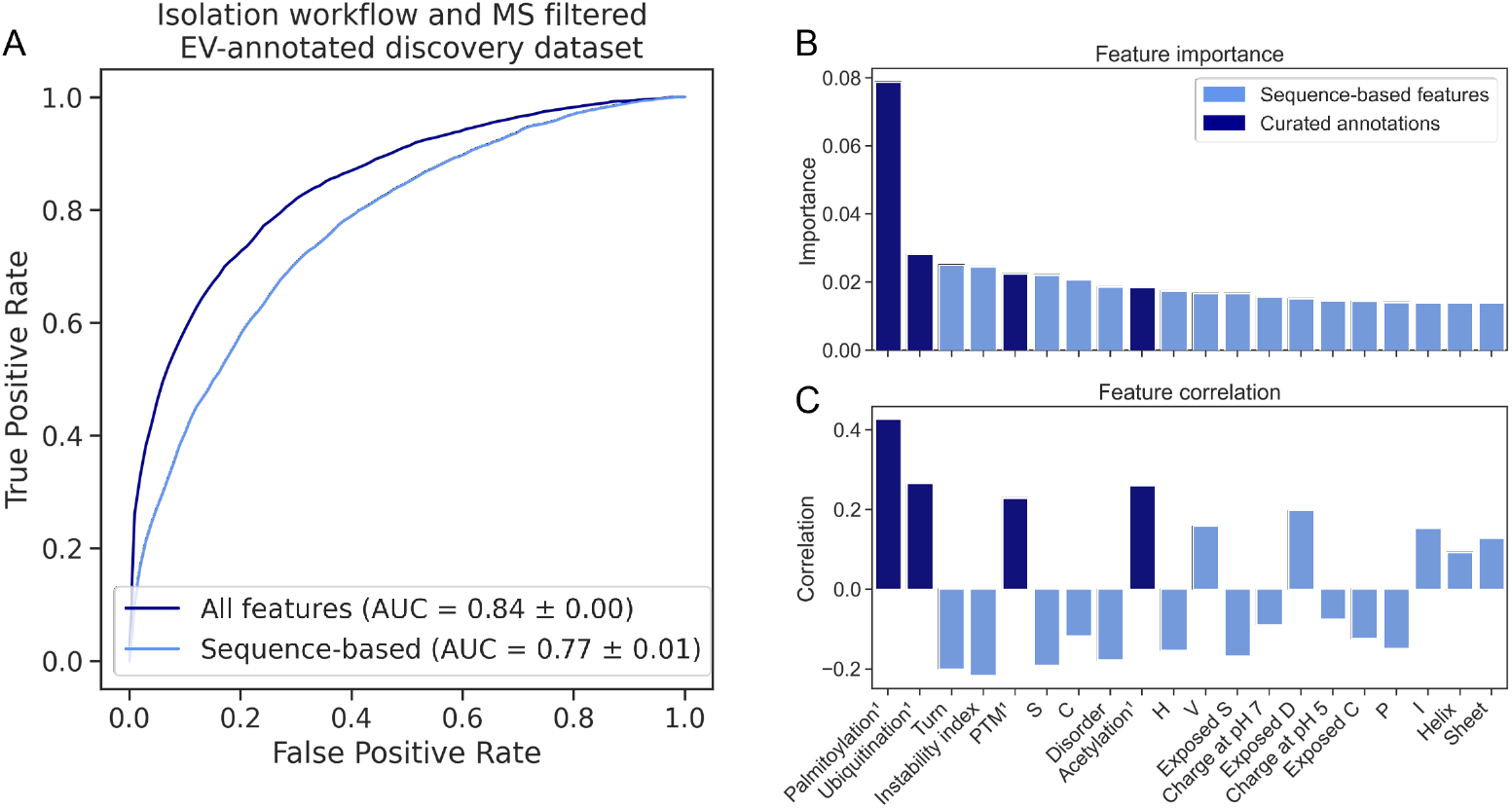
Performance of RF model and feature importance analysis. (**A**) ROC curves and AUC display the performance of the RF classifiers using sequence-based features (light blue) and sequence-based features and curated annotations (dark blue). (**B**) Bar plots show the Gini importance of the top 20 features for EV prediction (**A**) as well as the correlation of these features with the EV class (**C**). PTMs, stability, structure, and polarity differentiate EV and non-EV proteins. Dark blue features are curated annotations. AUC - area under the curve, C - cysteine, D - aspartic acid, EV - extracellular vesicle, H - histidine, I - isoleucine, P - proline, PTM - post-translational modification, RF - random forest, ROC - receiver operating characteristic, S - serine, V - valine

The model trained on the discovery set achieved a ROC-AUC of 0.77±0.01, which increased further to 0.84±0.00 when incorporating curated annotations as input. The superior model performance, when also including curated annotations, suggests that this type of information is valuable for the correct prediction of EV association and cannot be compensated for by sequence-based predictions alone.

We also trained models on the alternative datasets that excluded some of the filter steps (Figure S2). The slightly better performance of the classifier which is not trained on an MS-filtered dataset indicates that this classifier effectively (also) predicts MS detectability.

### Various features are important for prediction of EV association

To gain insight into the differences between EV and non-EV associated human proteins, we examined the protein features most decisive for the correct classification of the proteins. Figure 3B displays the Gini importance of the 20 most important features for EV association of the model trained on the discovery set. Figure 3C indicates for these 20 features if the correlation with the EV class is positive or negative. To compare which trends are identical and diverging in datasets with fewer filter steps, a heat map ranking all features across the three trained models was generated (Figure S3).

PTM annotations comprise evidently many of the most important features. Notably, all PTM types investigated here are positively correlated with the EV class indicating enrichment of PTMs in EV associated proteins. Especially palmitoylation shows high correlation (Figure 3C) and is by far the most important feature used by the model to predict EV association (Figure 3B). Sequence-based protein features considered important for prediction include predicted structure (Turn, Disorder, Helix, Sheet), stability (Instability index), and charge (Charge at pH 5 and 7, Exposed D) (Figure 3B). The abundances of less common lipidation PTMs, i.e., prenylation, myristoylation, and GPI-anchors, are shown Figure S4A. We were especially interested in those PTMs as palmitoylation, also a lipidation modification, was found to be so strongly enriched in the human EV proteome. Other rare PTMs have been linked to EV sorting [25, 27, 28]: citrullination, ISGylation, nitration and NEDDylation (Figure S4B). While for these PTM types the number of associated proteins is extremely limited, we could still show enrichment in the EV group. Less frequent PTMs were not important for EV classification by the RF because of their scarce absolute annotation numbers (Figure S3); nevertheless, these are biologically interesting.

The feature importance of *curated* PTM annotations was much higher than the importance of *predicted* PTMs by MusiteDeep. Hence, the differences between EV and non-EV proteins in terms of PTM annotations cannot be fully captured by state-of-the-art prediction methods that are purely sequence-based (Figure S3).

### Observations hold true in a high confidence EV set

In order to validate the determinants for EV association found by the machine learning models (Figure 3B) and simple correlation analysis (Figure 3C), we devised sets of EV annotations with different degrees of reliability.

We combined three recent studies from three different biological fluids which used effective isolation techniques, reducing incorrect EV annotations, *e.g*., due to lipoproteins or platelets [16], resulting in a set of *High Confidence* EV annotations. In addition, we devised a *Low Confidence* EV annotation set using older studies, for which we would expect contaminants to be included caused by insufficient isolation workflows.

Figure 4 shows the comparison between physicochemical and structural characteristics (Figure 4A-B), as well as PTM annotations (Figure 4C-D), of the high confidence validation EV proteins (HC EV), the proteins annotated by low confidence EV studies (LC EV), and the EV and non-EV datasets of the discovery set which was used for training and testing our prediction model. The associations of the investigated features that we identified in our EV dataset become even stronger in the high-confidence EV dataset, while identified EV-specific signals become diluted in the low-confidence EV set. All selected sequence-based features consistently show a clear trend: the difference in each feature between non-EV the EV sets increases, the higher confidence of the data (Figure 4A-B), suggesting that we identified true EV-specific determinants in our EV discovery dataset. The confirmation of the PTM properties of EV proteins is evident as well, especially for palmitoylation and nitrosylation (Figure 4C). In fact, in the high-confidence EV set, over half of the proteins contain a palmitoylation site, confirming the prevalence of this EV sorting mechanism. On the other hand, the PTM signals become diluted in the low-confidence dataset, probably due to many falsely reported discoveries in the experimental data. These results suggest which EV association determinants may be used for quality control purposes of experimental data.

**Figure 4.**
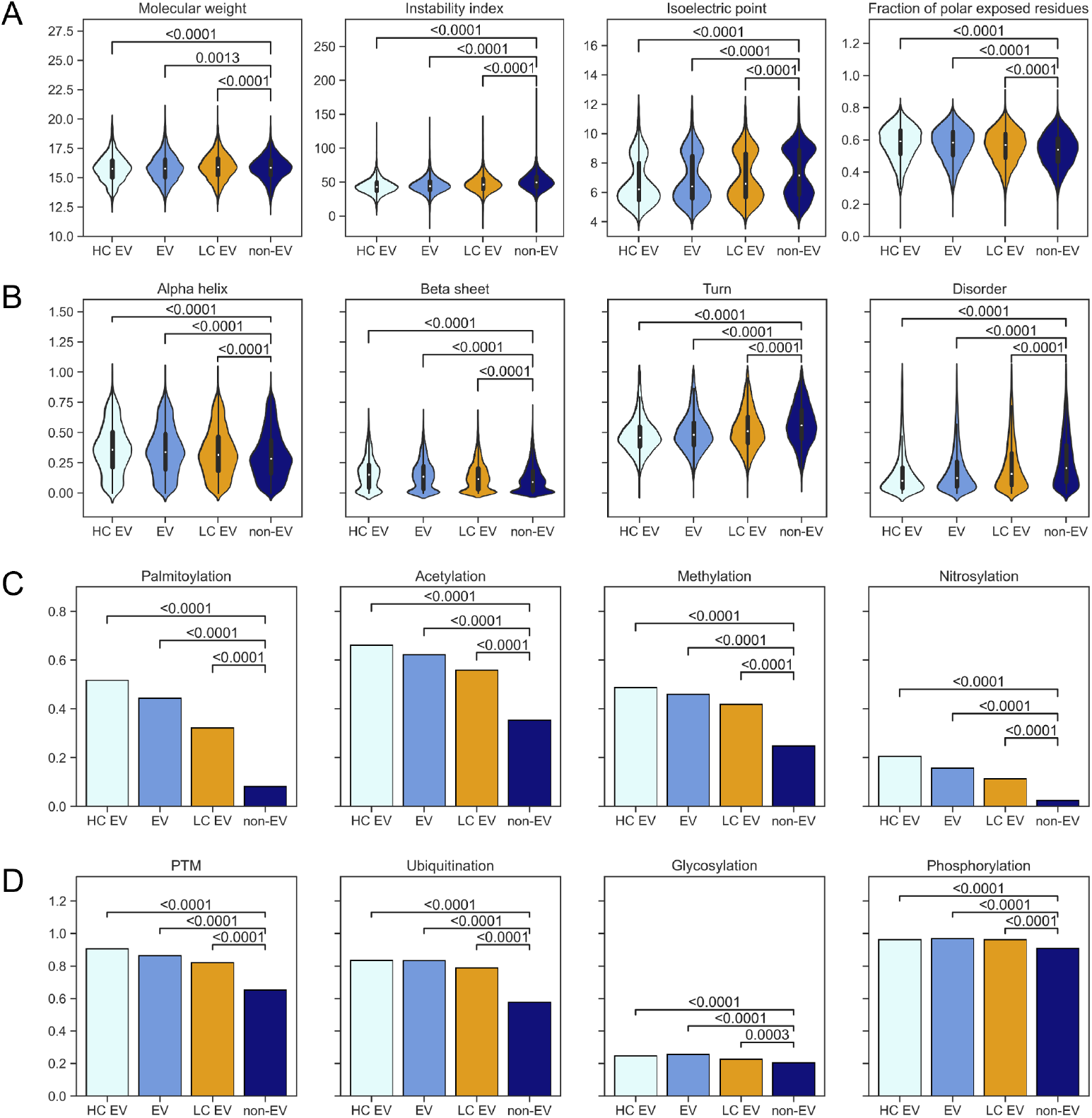
Features in the high and low confidence EV sets. Proteins in the high-confidence EV dataset, which was constructed from three recent studies show a similar distribution of physicochemical and structural properties as the EV protein set of the discovery dataset. For many features, the discrepancy with the non-EV group becomes more distinct. Furthermore, the low confidence dataset (orange) which contains EV proteins identified in older studies dilutes the observed signal compared to the EV protein set probably due to many falsely included contaminants. P-values are displayed in the plots. EV - extracellular vesicle, HC - high confidence, LC - low confidence.

### Functional characterisation of frequently detected EV proteins

In addition to the general characteristics of the EV proteome, we were interested in proteins that are often identified in EV studies and their properties. To functionally characterise EV associated proteins, we selected 478 unique human proteins from Vesiclepedia with occurrences (counts) in at least 30 different studies. Pathway enrichment analysis revealed proteins most frequently detected in EVs to be strongly associated with the ribosome. Notably, ribosomal proteins have been previously reported to be enriched in exosomes [29]. Other enriched pathways include the phagosome, cell-cell communication, and immune response pathways, including viral carcinogenesis, Escherichia coli infection, and antigen presentation (Figure S5).

### SHAP plots illustrate the model’s decision-making for case examples

After establishing the most important characteristics of EV proteins on a global level, we aimed to illustrate why the model predicts a protein to be EV associated (or not) for individual proteins. Note that this type of information cannot be derived directly for experimental annotations of EV association, and hence the prediction model can provide additional insight into possible sorting mechanisms associated with a specific protein.

Using SHAP values, the features that cause the model to predict EV association for a specific protein were analysed (Figure 5). SHAP plots show the importance of PTM presence for each case and their contribution to the predicted label.

**Figure 5.**
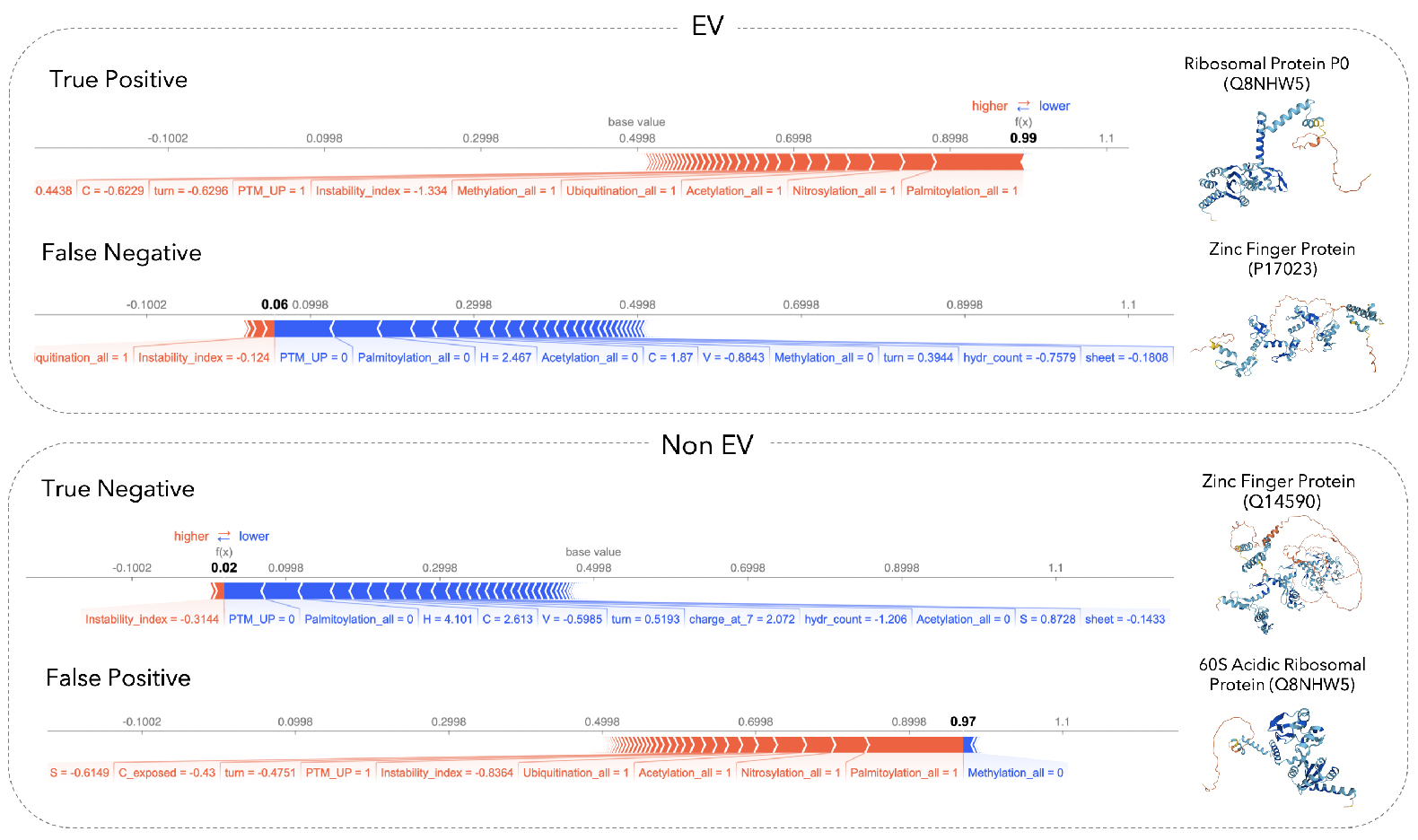
Shapley value analysis of case examples. SHAP plots for local interpretability are shown for correctly predicted proteins (i.e., true positive, true negative), as well as proteins for which the model prediction and the annotation from our data curation workflow do not agree with each other (i.e., false positive, false negative) from our test set. We chose examples in which the predictor is very certain if the individual protein is EV associated or not. Each SHAP plot displays a set of SHAP values that explain for each individual protein which features contributed to the model’s prediction. Features in red contribute to the prediction being higher (i.e., EV associated), and features in blue decrease the predicted score (i.e., non-EV).

Of interest is the case study of a false positively predicted protein, the ribosomal protein 60S acidic ribosomal protein P0-like (UniProt ID: Q8NHW5). Based on the properties this protein follows the general trend of EV proteome: it is heavily post-translationally modified and is stable, leading the model to predict the protein annotated as non-EV to be confidently classified as an EV associated protein. Investigation of this protein in our data revealed that while this protein is reported in multiple EV studies according to Vesiclepedia [17], its Gene (Entrez) ID could not be mapped to its UniProt entry. Thus, it was removed during the filtering steps despite being a true EV protein. Nevertheless, our model could correctly classify it.

## Discussion

In this work, we asked if it is possible to predict EV association based on the sequence (and associated annotations) of a protein, and which protein properties contributed to EV association predictions, both on a proteome- and an individual protein level.

An extensive multi-step data curation and filtering workflow allowed us to discriminate true determinants of EV association from an experimental bias inherent to EV databases. Importantly, we analysed unique proteins regarding their presence or absence in EVs, for which defining clear positives and negatives is essential. We showed that it is indeed possible to predict EV association from amino acid sequences with an AUC of 0.77 in our filtered discovery set (Figure 3A). The performance increased further to 0.84 when incorporating curated annotations. Feature analysis of the EV and non-EV proteome elucidated that EV proteins are more stable, polar, and structured than non-EV proteins and contain various PTM sites (Figure 3B-C). These observed biological trends held true in a high-confidence EV set of recent, high-quality EV studies (Figure 4), confirming the determinants of EV association. Furthermore, we show that the low-confidence dataset containing proteins identified in older EV studies mostly weakens the signal (Figure 4) highlighting the importance of data curation steps suggested in this work.

We showed that experimental MS data of EV studies are biased towards larger proteins, and that correction during the data curation process is required. The presented approach to correct protein annotations for the introduced bias of MS is broadly applicable to other proteomics-based studies and databases. As many, especially older studies did not contemplate any standardization or possible contaminants, spurious annotations are highly likely to be present in EV databases. This bias had to be counteracted by limiting the studies to be included and excluding low-count proteins. Note that because this study only considered the presence or absence of a protein in EVs, the impact of a few contaminating proteins that were not successfully excluded from the EV set is limited.

While the accuracy of the simple RF classifier is encouraging, improved performance is conceivable, *e.g*., by the incorporation of residue-specific instead of solely global features [30, 31]. As of yet, there has been no attempt to apply deep learning approaches to this task which is a promising approach to pursue but might be limited by the amount of available training data. Notably, this study suggests that smaller, but stricter datasets lead to more trustworthy classifiers.

Several of our findings about EV properties correspond to already established knowledge about these vesicles. While in both EV and non-EV proteins the typical bimodal distribution of the isoelectric point was present [32], an enrichment of proteins with a lower isoelectric point, *i.e*., of more acidic proteins, in the EV proteome was found in our analysis (Figure 3C, Figure 4). Interestingly, previous work on exosomes has shown that acidic conditions favor the existence and release of those vesicles [33,34]. Another possible explanation for this preference towards acidity is the relationship between isoelectric point distribution and subcellular localization. The proteomes of the cytoplasm, lysosome, vacuoles, and cytoskeleton have been shown to have an acidic protein composition, while proteins of the plasma membrane and mitochondria are more basic [32, 35]. Our findings could indicate an enrichment of the former subcellular proteomes, and a depletion of the latter within EVs. An overlap of EV and lysosome proteomes is particularly expected as the sorting of proteins into both exosomes and lysosomes is dependent on multivesicular bodies (MVBs) [36]. An enrichment of cytoplasmic proteins is also coherent as the biogenesis of both exosomes and microvesicles involves the envelopment and budding of cytosolic components at the MVB or plasma membrane, respectively [37].

The importance of PTMs in the biogenesis, cargo-loading, and release of EVs has been studied intensively [25, 38, 39]. Our work supports previous findings about the existence or even abundance of phosphorylation [40], glycosylation [41, 42], and ubiquitination [38, 43, 44] in EV proteins. We also detected a higher fraction of proteins with methylation and acetylation sites despite no previous strong evidence of their relevance for EV sorting. However, these modifications might not be specific to the EV protein sorting process as their correlation became weaker with an increasing number of filtering steps (Figure S3).

Nitrosylation has been linked to synaptic vesicles [45, 46] but its general function remains elusive as it has been implicated in cell survival and death, regulation of protein activity, and localization amongst others [47, 48]. As a clear enrichment of nitrosylation sites was found in the EV proteome (Figure 4), their potential role in protein localization into EVs should be explored. Palmitoylation emerged to be the strongest feature for EV association - with over half of the high confidence EV associated proteins containing a palmitoylation site Figure 4. Interestingly, palmitoylation has been implicated in recent years in the EV sorting of specific proteins [49, 50]. Moreover, Mariscal *et al*. showed in a comprehensive study of the EV palmitoyl-proteome a high abundance of palmitoylated proteins in cancer-derived EVs [51]. The enrichment in the human EV proteome for both palmitoylation, as well as other types of lipidation, may be explained by their involvement in binding proteins to the plasma membrane [52].

The functional enrichment analysis results for the most frequently detected EV proteins are in line with what is currently considered as major functions of EVs, namely cell-to-cell communication [36, 53], signalling via membrane receptors or cell fusion by releasing cargo [3]. Moreover, evidence for an immunoregulatory function of EVs is increasing [54] as exosomes were shown to stimulate immune responses by presenting antigens on their surface [55] or as carriers of tumor and pathogenic antigens [56–58]. Furthermore, a recent investigation of EVs shed light on their involvement in various disease pathologies including metabolic disorders [59] and viral infection [60].

Taken together, our study provides a systematic characterisation of the EV proteome and a machine learning tool for predicting EV association. Moreover, we illustrate how to provide a readout of features for specific proteins. Based on our machine learning approach, we highlight possible protein sorting mechanisms and quality control criteria that can guide future EV experiments and downstream analysis.

## Methods

A data curation workflow incorporating several resources and filtering steps was created to obtain protein sets annotated regarding their presence or absence in EVs (Figure 1). We accessed online databases to identify human proteins associated with EVs. Simultaneously, we generated a dataset of all unique human proteins with various sequence-based features, as well as curated annotations. Several different filtering strategies were applied to receive higher confidence EV and non-EV protein sets. The resulting EV and non-EV annotations were used to train and test machine learning classifiers and analyse the features necessary for the correct classification of EV association.

### Dataset generation

#### EV dataset

The EV proteome was downloaded from the EV databases Vesiclepedia (http://microvesicles.org, Version 4.1) [17] and ExoCarta (http://www.exocarta.org, Version 5) [18, 19]. Entries were filtered for human protein content, and the ‘counts’, *i.e*., the number of experiments that identified each protein, were calculated. After merging the entries from both databases, 13,648 unique proteins were identified as EV associated (Figure 1). For 11,952 of these proteins the unique UniProt ID could be determined which comprise the *General EV dataset*.

#### Human proteome dataset

Swiss-Prot reviewed human proteins were extracted from UniProt [61] (release: 2020 05) containing 20,385 unique proteins. For 20,381 of these proteins, the full feature set could be generated which comprises the *Human proteome* dataset.

#### Filtering steps for the EV-annotated discovery dataset

To generate a model able to predict which proteins are associated with EVs, it is essential to have a training dataset that is as accurately labelled as possible. The steps below describe, how we filtered and annotated our datasets to minimise the experimental and publication bias, resulting in our *discovery dataset*.

Firstly, we generated an *isolation method filter* (Figure 1). The filter excludes proteins that were identified through workflows prone to a high false discovery rate. The information provided on Vesiclepedia regarding applied isolation methods was used to include solely proteins identified in experiments that: 1. made no use of ExoQuick or similar “high recovery but low purity” isolation kits; 2. were comprised of at least three different isolation steps that enrich for different properties (e.g., size and density); 3. did not apply ultracentrifugation as the first step in their EV isolation workflow. This approach resulted in 6,818 retained EV proteins of which 6,678 were mapped to respective UniProt IDs.

Secondly, we created a *MS filter* (see Figure 1). As MS was predominantly utilized for protein detection in the majority of EV studies, experimental detection bias known to be introduced by MS [26] is likely to exist in this data. This is specifically problematic for the proteins we label as non-EV associated. Note that these proteins may simply not have been detected in EV studies because of the limitations of MS as a detection method. Thus, we wished to exclude any proteins that have never been identified in MS experiments from the discovery dataset to counteract potential bias. For this purpose, we accessed ProteomicsDB, an expansive collection of MS studies of the human proteome among others [62, 63]. We collected all human proteins in this database that have been verifiably detected in previous MS studies. Only proteins that are present in both the ProteomicsDB and the human proteome datasets were retained for further analysis (16,790 proteins). By excluding any entries from the human proteome dataset not found in proteomics studies an *MS-detectable human proteome* dataset was generated (Figure 1).

Finally, we removed low-count proteins, i.e., those found in less than three unique experiments, from the entire dataset as those proteins were considered ambiguous. Thus, these are neither included as EV nor non-EV proteins.

The final discovery dataset contains 5,965 and 10,290 EV and non-EV annotated proteins, respectively. To examine the effect of the filtering steps described above, we also trained models based on datasets that skip some of the filtering steps, the workflow for generating these alternative discovery sets is provided in Figure S1.

### Feature generation

We generated a range of features to identify properties that significantly differentiate between proteins identified in EVs and those never detected in EVs. In total, 95 features were included for each protein present in our datasets. In order to obtain explainable machine learning models, it is essential to use input features that are easy to interpret, and that can help to reveal potential EV sorting mechanisms, as well as potential biases in the datasets. For this reason, a wide variety of interpretable protein properties was derived directly from the protein sequences; in addition, curated database annotations were added.

#### Sequence-based features

Table S1 lists all these features and details their generation.

Basic features, such as sequence length, molecular weight, number of residues per amino acid, and polar and hydrophobic amino acid count were calculated directly from the sequence. All amino acid counts were normalized by the length of the protein to obtain amino acid proportions. Sequence length and molecular weight were log2-transformed.

NetSurfP-2.0 [30] was used to predict solvent accessibility, secondary structure, and structural disorder for each residue in the protein sequences. The per-residue predictions were then utilized to calculate global features for each protein according to van Gils *et al*. [64]. Residues with relative surface accessibility (RSA) > 0.4 were considered exposed. The number of exposed residues per protein, for each amino acid, polar amino acids, and hydrophobic amino acids, were calculated and divided by the sum of all exposed amino acids of the respective protein. We calculated the proportion of structural elements (*α* helices, *β* sheets, turns, disordered regions), total accessible surface area (TASA), and total hydrophobic surface area (THSA). TASA and THSA were log2-transformed. Relative hydrophobic surface area (RHSA) was derived as the fraction of THSA relative to TASA.

The Bio.SeqUtils.ProtParam Biopython module [65] was used to calculate aromaticity [66], instability index [67], Gravy [68], isoelectric point, charge at pH-7 and pH-5. The SoDoPe tool [69] was used to derive the probability of solubility. A global aggregation propensity score was calculated for each protein. The aggregation propensity score of each amino acid was derived from the experimental work by De Groot *et al*. [70].

For various domains, specifically coiled-coil, WW domains, RAS profiles, EGF and RRM, the associated PROSITE [71–73] identifiers were extracted (see Table S1). ScanProsite [74] was applied to cross-reference protein sequences and domain motifs against each other and the domain presences were implemented as binary features.

To enable the use of PTMs as a sequence-based feature, MusiteDeep PTM prediction results were incorporated as features [75, 76]. A threshold of 0.75 was chosen to include a predicted PTM site and only the highest-scoring PTM was considered at every site. The predicted PTM sites were implemented as a global feature for each protein, indicating if a PTM is present in or absent from the protein sequence.

TMHMM [77] was used to predict transmembrane helices that were included as a binary feature indicating presence or absence.

#### Curated annotations

The databases used to create curated annotation and relevant comments are listed in Table S2.

Databases of PTM annotations were accessed to annotate the human proteome. Relevant datasets were downloaded from PhosphoSitePlus [78], iPTMnet [79], Swiss-Palm [80, 81] and UniProtKB [61]. Annotations for two uncommon PTMs (ISGylation, NEDDylation) were extracted from UniProt via text mining of the comments section of sequence position independent annotations. While some of these databases also provided position-specific modification information, all feature annotations were generated at a protein level indicating if at least one amino acid in the protein sequence is known to be modified regarding each PTM type. UniProt annotations for transmembrane and heat-shock proteins were administered as binary features.

### Prediction of EV association

To ascertain the possibility to predict the EV association of a protein from the sequence, as well as to determine which features are considered most important for this task, random forest (RF) classification was implemented. The three discovery datasets described in the section *Datasets* were used and for each of those datasets, two RF models were trained: using only sequence-based features and using sequence-based features and curated annotations. This resulted in six classifiers.

#### Training and interpretation of the random forest

The discovery dataset was split into an 80% training and 20% testing set, resulting in 9,544 training and 3,251 testing entries. As the classes (EV and non-EV) were unbalanced, undersampling of the class containing a higher number of proteins, *i.e*., the majority class, was utilized to create balanced training datasets. All continuous features were scaled using the robust scaler provided in the `scikit-learn` Python library [82].

The `scikit-learn` RF model was implemented. To compare the importance of the features for the prediction of EV association the impurity-based feature importance of the RF models, *i.e*., the Gini importance, was assessed. Higher values indicate greater importance for correct classification. Model performance was evaluated using the ROC curve and the AUC score. To illustrate specific cases, Shapley Additive explanations (SHAP) analysis was carried out [83]. ‘Feature interactions’ were calculated to highlight feature combinations used for predictions.

#### Validation of EV association determinants

We aimed to validate the protein-encoded determinants that were found to be important features in the machine learning models, by observing trends of these properties in high- and low-confidence EV datasets in terms of experimental procedures.

Since present-day EV studies are characterised by superior enrichment and isolation methods, we chose three recent EV studies of different body fluids that have not been included on Vesiclepedia or ExoCarta yet: 1,789 proteins analysed from urine EVs [84], 1,187 proteins from plasma EVs [16], and 1,686 proteins from breast cancer cell line MDA-MB-468 cell culture media EVs [85]. We combined the identified EV associated proteins into a high-confidence EV dataset containing 3,222 proteins, out of which 573 were not annotated as EV proteins in our discovery dataset. Secondly, we constructed a low-confidence dataset by only selecting proteins from Vesiclepedia identified in EV studies published before the year 2010. Out of 9,506 unique proteins, 4,097 were not annotated as EV proteins in our discovery set.

We hypothesized that the high-confidence dataset will display the same trends of protein properties as the EV proteins of our discovery set; the existence of non-EV contaminants should be limited because of the high isolation standards of the included studies. We expected the signal in the low-confidence dataset to be diluted by contaminants, demonstrating the missing specificity of early EV isolation workflows.

### Statistical analysis

To test for the significance of features, we employed Mann-Whitney U test and Fisher’s exact test for continuous and categorical features, respectively. The threshold for significance was set at .05 and correction for multiple testing was done using the Benjamini-Hochberg procedure.

### Functional analysis of EV associated proteins

To analyse proteins that are often identified in EVs (top EV proteins), 478 unique human proteins from Vesiclepedia with occurrences (counts) in at least 30 different studies were selected from our discovery set and analysed. GSEApy enrichr tool was used for the pathway enrichment analysis using the KEGG 2019 Human library [86].

## Data availability

All code and data related to this work can be found on: https://github.com/ibivu/ExtracellularVesicles

## Acknowledgements

We would like to thank Jose Gavaldá-García for exploring the possibility of this study.

## Supporting Information

**Figure S1**

**Data curation workflow for less stringently filtered datasets**. Two alternative datasets were generated by a similar data curation workflow as described for the discovery set albeit some filtering steps were skipped. The general EV dataset was used to annotate the human proteome to obtain the study count filtered set (red). The MS-detectable human proteome was annotated by the general EV dataset leading to the study count and MS-filtered set (green). Both datasets are less stringently filtered than the discovery set (Figure 1) as no isolation workflow filtering was performed. EV - extracellular vesicle, MS - mass spectrometry.)

**Figure S2**

**Performance of RF models trained on less stringently filtered datasets**. 470 RF models trained on datasets that skipped some of the filtering steps show similar performance for EV association prediction. The unfiltered dataset (red) performs slightly better indicating that this classifier effectively (also) predicts MS detectability. Including the isolation workflow filter does not affect the model performance significantly.

**Figure S3**

**Heat map of feature importance ranking**. The feature importances of the three RF models are ranked. The RFs have been trained on datasets of differing stringency, i.e., unfiltered (red), MS-filtered (green) and isolation and MS-filtered datasets. Marked features (^1^) are curated annotations. MS - mass spectrometry, RF - random forest.

**Figure S4**

**Uncommon PTM annotations across the EV and non-EV proteome**. The fraction of annotated proteins regarding less common PTM types show stark differences between EV and non-EV classes. All included PTM types are enriched in the EV class but the absolute number of annotations is scarce. EV - extracellular vesicle, PTM - post-translational modification.

**Figure S5**

**Enrichment analysis of the most common EV proteins**. Proteins that are often identified in EVs (top EV proteins with occurrences in at least 30 different studies) were selected from our discovery set III and analysed with GSEApy enrichr tool (KEGG 2019 Human library) [86]. KEGG pathways that are enriched include ribosome, phagosome, proteasome, and pathways associated with infection and immune response.

**Table S1**

Description of all generated sequence-based features implemented in the random forest classifiers.

**Table S2**

Description of all annotation features implemented in the random forest classifiers.

## References

1. F. T. Borges, L. Reis, and N. Schor, “Extracellular vesicles: structure, function, and potential clinical uses in renal diseases,” Brazilian Journal of Medical and Biological Research, vol. 46, no. 10, pp. 824–830, 2013.

2. M. Yáñez-Mó, P. R.-M. Siljander, Z. Andreu, A. Bedina Zavec, F. E. Borrás, E. I. Buzas, K. Buzas, E. Casal, F. Cappello, J. Carvalho, et al., “Biological properties of extracellular vesicles and their physiological functions,” Journal of extracellular vesicles, vol. 4, no. 1, p. 27066, 2015.

3. G. van Niel, G. D’Angelo, and G. Raposo, “Shedding light on the cell biology of extracellular vesicles,” Nature Reviews Molecular Cell Biology, vol. 19, pp. 213–228, Jan. 2018.

4. A. Gámez-Valero, K. Beyer, and F. E. Borràs, “Extracellular vesicles, new actors in the search for biomarkers of dementias,” Neurobiology of Aging, vol. 74, pp. 15–20, Feb. 2019.

5. K. C. Miranda, D. T. Bond, M. McKee, J. Skog, T. G. Păunescu, N. Da Silva, D. Brown, and L. M. Russo, “Nucleic acids within urinary exosomes/microvesicles are potential biomarkers for renal disease,” Kidney international, vol. 78, no. 2, pp. 191–199, 2010.

6. T. L. Whiteside, “The potential of tumor-derived exosomes for noninvasive cancer monitoring,” Expert review of molecular diagnostics, vol. 15, no. 10, pp. 1293–1310, 2015.

7. T. Camino, N. Lago-Baameiro, S. B. Bravo, A. Molares-Vila, A. Sueiro, I. Couto, J. Baltar, E. F. Casanueva, and M. Pardo, “Human obese white adipose tissue sheds depot-specific extracellular vesicles and reveals candidate biomarkers for monitoring obesity and its comorbidities,” Translational Research, 2021.

8. L. S. Watson, E. D. Hamlett, T. D. Stone, and C. Sims-Robinson, “Neuronally derived extracellular vesicles: An emerging tool for understanding alzheimer’s disease,” Molecular Neurodegeneration, vol. 14, 6 2019.

9. J. Lötvall, A. F. Hill, F. Hochberg, E. I. Buzás, D. D. Vizio, C. Gardiner, Y. S. Gho, I. V. Kurochkin, S. Mathivanan, P. Quesenberry, S. Sahoo, H. Tahara, M. H. Wauben, K. W. Witwer, and C. Théry, “Minimal experimental requirements for definition of extracellular vesicles and their functions: a position statement from the international society for extracellular vesicles,” Journal of Extracellular Vesicles, vol. 3, p. 26913, Jan. 2014.

10. C. Théry, K. W. Witwer, E. Aikawa, M. J. Alcaraz, J. D. Anderson, R. Andriantsi-tohaina, A. Antoniou, T. Arab, F. Archer, G. K. Atkin-Smith, et al., “Minimal information for studies of extracellular vesicles 2018 (misev2018): a position statement of the international society for extracellular vesicles and update of the misev2014 guidelines,” Journal of extracellular vesicles, vol. 7, no. 1, p. 1535750, 2018.

11. S. Gandham, X. Su, J. Wood, A. L. Nocera, S. C. Alli, L. Milane, A. Zimmerman, M. Amiji, and A. R. Ivanov, “Technologies and standardization in research on extracellular vesicles,” Trends in Biotechnology, vol. 38, pp. 1066–1098, Oct. 2020.

12. K. W. Witwer, E. I. Buzás, L. T. Bemis, A. Bora, C. Lässer, J. Lötvall, E. N. N. ‘t Hoen, M. G. Piper, S. Sivaraman, J. Skog, C. Théry, M. H. Wauben, and F. Hochberg, “Standardization of sample collection, isolation and analysis methods in extracellular vesicle research,” Journal of Extracellular Vesicles, vol. 2, p. 20360, jan 2013.

13. F. A. Coumans, A. R. Brisson, E. I. Buzas, F. Dignat-George, E. E. Drees, S. El-Andaloussi, C. Emanueli, A. Gasecka, A. Hendrix, A. F. Hill, et al., “Methodological guidelines to study extracellular vesicles,” Circulation research, vol. 120, no. 10, pp. 1632–1648, 2017.

14. J. L. Welton, J. P. Webber, L.-A. Botos, M. Jones, and A. Clayton, “Ready-made chromatography columns for extracellular vesicle isolation from plasma,” Journal of extracellular vesicles, vol. 4, no. 1, p. 27269, 2015.

15. B. W. Sódar, Á. Kittel, K. Pálóczi, K. V. Vukman, X. Osteikoetxea, K. Szabó-Taylor, A. Németh, B. Sperlágh, T. Baranyai, Z. Giricz, et al., “Low-density lipoprotein mimics blood plasma-derived exosomes and microvesicles during isolation and detection,” Scientific reports, vol. 6, no. 1, pp. 1–12, 2016.

16. N. Karimi, A. Cvjetkovic, S. C. Jang, R. Crescitelli, M. A. H. Feizi, R. Nieuwland, J. Lötvall, and C. Lässer, “Detailed analysis of the plasma extracellular vesicle proteome after separation from lipoproteins,” Cellular and molecular life sciences, vol. 75, no. 15, pp. 2873–2886, 2018.

17. M. Pathan, P. Fonseka, S. V. Chitti, T. Kang, R. Sanwlani, J. V. Deun, A. Hendrix, and S. Mathivanan, “Vesiclepedia 2019: a compendium of RNA, proteins, lipids and metabolites in extracellular vesicles,” Nucleic Acids Research, vol. 47, pp. D516–D519, Nov. 2018.

18. R. J. Simpson, H. Kalra, and S. Mathivanan, “Exocarta as a resource for exosomal research,” Journal of extracellular vesicles, vol. 1, no. 1, p. 18374, 2012.

19. S. Keerthikumar, D. Chisanga, D. Ariyaratne, H. Al Saffar, S. Anand, K. Zhao, M. Samuel, M. Pathan, M. Jois, N. Chilamkurti, et al., “Exocarta: a web-based compendium of exosomal cargo,” Journal of molecular biology, vol. 428, no. 4, pp. 688–692, 2016.

20. K. Waury, E. A. J. Willemse, E. Vanmechelen, H. Zetterberg, C. E. Teunissen, and S. Abeln, “Bioinformatics tools and data resources for assay development of fluid protein biomarkers,” Biomarker Research, vol. 10, nov 2022.

21. R. Kumar and S. K. Dhanda, “Bird eye view of protein subcellular localization prediction,” Life, vol. 10, no. 12, p. 347, 2020.

22. L. Zhao, G. Poschmann, D. Waldera-Lupa, N. Rafiee, M. Kollmann, and K. Stühler, “Outcyte: A novel tool for predicting unconventional protein secretion,” Scientific reports, vol. 9, no. 1, pp. 1–9, 2019.

23. B. Liu, L. Leng, X. Sun, Y. Wang, J. Ma, and Y. Zhu, “Ecmpride: prediction of human extracellular matrix proteins based on the ideal dataset using hybrid features with domain evidence,” PeerJ, vol. 8, p. e9066, 2020.

24. A. Ras-Carmona, M. Gomez-Perosanz, and P. A. Reche, “Prediction of unconventional protein secretion by exosomes,” BMC Bioinformatics, vol. 22, jun 2021.

25. S. Anand, M. Samuel, S. Kumar, and S. Mathivanan, “Ticket to a bubble ride: Cargo sorting into exosomes and extracellular vesicles,” Biochimica et Biophysica Acta (BBA)-Proteins and Proteomics, vol. 1867, no. 12, p. 140203, 2019.

26. F. Klont, L. Bras, J. C. Wolters, S. Ongay, R. Bischoff, G. B. Halmos, and P. Horvatovich, “Assessment of sample preparation bias in mass spectrometry-based proteomics,” Analytical Chemistry, vol. 90, pp. 5405–5413, Apr. 2018.

27. O. Moreno-Gonzalo, C. Villarroya-Beltri, and F. SÃ¡nchez-Madrid, “Post-translational modifications of exosomal proteins,” Frontiers in Immunology, vol. 5, Aug. 2014.

28. C. Campanella, A. D’Anneo, A. M. Gammazza, C. C. Bavisotto, R. Barone, S. Emanuele, F. L. Cascio, E. Mocciaro, S. Fais, E. C. D. Macario, A. J. Macario, F. Cappello, and M. Lauricella, “The histone deacetylase inhibitor SAHA induces HSP60 nitration and its extracellular release by exosomal vesicles in human lung-derived carcinoma cells,” Oncotarget, vol. 7, pp. 28849–28867, Dec. 2015.

29. V. Dozio and J.-C. Sanchez, “Characterisation of extracellular vesicle-subsets derived from brain endothelial cells and analysis of their protein cargo modulation after tnf exposure,” Journal of extracellular vesicles, vol. 6, no. 1, p. 1302705, 2017.

30. M. S. Klausen, M. C. Jespersen, H. Nielsen, K. K. Jensen, V. I. Jurtz, C. K. Sønderby, M. O. A. Sommer, O. Winther, M. Nielsen, and B. Petersen, “Netsurfp-2.0: Improved prediction of protein structural features by integrated deep learning,” Proteins: Structure, Function, and Bioinformatics, vol. 87, pp. 520–527, 2019.

31. S. M. A. Shah and Y.-Y. Ou, “Trp-bert: Discrimination of transient receptor potential (trp) channels using contextual representations from deep bidirectional transformer based on bert,” Computers in Biology and Medicine, vol. 137, p. 104821, 2021.

32. J. Kiraga, P. Mackiewicz, D. Mackiewicz, M. Kowalczuk, P. Biecek, N. Polak, K. Smolarczyk, M. R. Dudek, and S. Cebrat, “The relationships between the isoelectric point and: Length of proteins, taxonomy and ecology of organisms,” BMC Genomics, vol. 8, 6 2007.

33. I. Parolini, C. Federici, C. Raggi, L. Lugini, S. Palleschi, A. D. Milito, C. Coscia, E. Iessi, M. Logozzi, A. Molinari, M. Colone, M. Tatti, M. Sargiacomo, and S. Fais, “Microenvironmental pH is a key factor for exosome traffic in tumor cells,” Journal of Biological Chemistry, vol. 284, pp. 34211–34222, Dec. 2009.

34. J.-J. Ban, M. Lee, W. Im, and M. Kim, “Low pH increases the yield of exosome isolation,” Biochemical and Biophysical Research Communications, vol. 461, pp. 76–79, May 2015.

35. A. Kurotani, A. A. Tokmakov, K.-I. Sato, V. E. Stefanov, Y. Yamada, and T. Sakurai, “Localization-specific distributions of protein pI in human proteome are governed by local pH and membrane charge,” BMC Molecular and Cell Biology, vol. 20, Aug. 2019.

36. H. Kalra, G. P. Drummen, and S. Mathivanan, “Focus on extracellular vesicles: introducing the next small big thing,” International journal of molecular sciences, vol. 17, no. 2, p. 170, 2016.

37. M. Mathieu, L. Martin-Jaular, G. Lavieu, and C. Théry, “Specificities of secretion and uptake of exosomes and other extracellular vesicles for cell-to-cell communication,” Nature Cell Biology, vol. 21, pp. 9–17, Jan. 2019.

38. H. Ageta and K. Tsuchida, “Post-translational modification and protein sorting to small extracellular vesicles including exosomes by ubiquitin and ubls,” Cellular and Molecular Life Sciences, vol. 76, pp. 4829–4848, 12 2019.

39. J. M. Carnino, K. Ni, and Y. Jin, “Post-translational modification regulates formation and cargo-loading of extracellular vesicles,” Frontiers in Immunology, vol. 11, May 2020.

40. P. A. Gonzales, T. Pisitkun, J. D. Hoffert, D. Tchapyjnikov, R. A. Star, R. Kleta, N. S. Wang, and M. A. Knepper, “Large-scale proteomics and phosphoproteomics of urinary exosomes,” Journal of the American Society of Nephrology, vol. 20, pp. 363–379, Dec. 2008.

41. J. Q. Gerlach and M. D. Griffin, “Getting to know the extracellular vesicle glycome,” Molecular BioSystems, vol. 12, pp. 1071–1081, 2016.

42. J. M. da Silva, V. F. Santiago, L. Rosa-Fernandes, C. R. Marinho, and G. Palmisano, “Protein glycosylation in extracellular vesicles: Structural characterization and biological functions,” Molecular Immunology, vol. 135, pp. 226–246, 7 2021.

43. S. I. Buschow, J. M. Liefhebber, R. Wubbolts, and W. Stoorvogel, “Exosomes contain ubiquitinated proteins,” Blood Cells, Molecules, and Diseases, vol. 35, pp. 398–403, Nov. 2005.

44. V. L. Smith, L. Jackson, and J. S. Schorey, “Ubiquitination as a mechanism to transport soluble mycobacterial and eukaryotic proteins to exosomes,” The Journal of Immunology, vol. 195, pp. 2722–2730, Aug. 2015.

45. Y. Wang, Z. Zhou, T. Leylek, H. Tan, Y. Sun, F. Parkinson, and J.-F. Wang, “Protein cysteine s-nitrosylation inhibits vesicular uptake of neurotransmitters,” Neuroscience, vol. 311, pp. 374–381, Dec. 2015.

46. Y. Wang, Z. Zhou, H. Tan, S. Zhu, Y. Wang, Y. Sun, X.-M. Li, and J.-F. Wang, “Nitrosylation of vesicular transporters in brain of amyloid precursor protein/presenilin 1 double transgenic mice,” Journal of Alzheimer’s Disease, vol. 55, p. 1683–1692, Dec. 2016.

47. Y. Koriyama and A. Furukawa, “S-nitrosylation regulates cell survival and death in the central nervous system,” Neurochemical Research, vol. 43, pp. 50–58, Jan. 2018.

48. C. T. Stomberski, D. T. Hess, and J. S. Stamler, “Protein s-nitrosylation: Determinants of specificity and enzymatic regulation of s-nitrosothiol-based signaling,” Antioxidants & Redox Signaling, vol. 30, pp. 1331–1351, Apr. 2019.

49. D. P. Romancino, V. Buffa, S. Caruso, I. Ferrara, S. Raccosta, A. Notaro, Y. Campos, R. Noto, V. Martorana, A. Cupane, A. Giallongo, A. d’Azzo, M. Manno, and A. Bongiovanni, “Palmitoylation is a post-translational modification of alix regulating the membrane organization of exosome-like small extracellular vesicles,” Biochimica et Biophysica Acta (BBA) - General Subjects, vol. 1862, pp. 2879–2887, Dec. 2018.

50. J. P. Flemming, B. L. Hill, M. W. Haque, J. Raad, C. S. Bonder, L. A. Harshyne, U. Rodeck, A. Luginbuhl, J. K. Wahl, K. Y. Tsai, P. J. Wermuth, A. M. Overmiller, and M. G. Mahoney, “miRNA- and cytokine-associated extracellular vesicles mediate squamous cell carcinomas,” Journal of Extracellular Vesicles, vol. 9, p. 1790159, July 2020.

51. J. Mariscal, T. Vagner, M. Kim, B. Zhou, A. Chin, M. Zandian, M. R. Freeman, S. You, A. Zijlstra, W. Yang, and D. D. Vizio, “Comprehensive palmitoyl-proteomic analysis identifies distinct protein signatures for large and small cancer-derived extracellular vesicles,” Journal of Extracellular Vesicles, vol. 9, p. 1764192, June 2020.

52. S. Picciotto, D. P. Romancino, V. Buffa, A. Cusimano, A. Bongiovanni, and G. Adamo, “Post-translational lipidation in extracellular vesicles: chemical mechanisms, biological functions and applications,” in Advances in Biomembranes and Lipid Self-Assembly, pp. 83–111, Elsevier, 2020.

53. R. C. Paolicelli, G. Bergamini, and L. Rajendran, “Cell-to-cell communication by extracellular vesicles: focus on microglia,” Neuroscience, vol. 405, pp. 148–157, 2019.

54. P. D. Robbins and A. E. Morelli, “Regulation of immune responses by extracellular vesicles,” Nature Reviews Immunology, vol. 14, pp. 195–208, Feb. 2014.

55. A. Bobrie, M. Colombo, G. Raposo, and C. Théry, “Exosome secretion: molecular mechanisms and roles in immune responses,” Traffic, vol. 12, no. 12, pp. 1659–1668, 2011.

56. J. Wolfers, A. Lozier, G. Raposo, A. Regnault, C. Théry, C. Masurier, C. Flament, S. Pouzieux, F. Faure, T. Tursz, E. Angevin, S. Amigorena, and L. Zitvogel, “Tumor-derived exosomes are a source of shared tumor rejection antigens for CTL cross-priming,” Nature Medicine, vol. 7, pp. 297–303, Mar. 2001.

57. J. D. Walker, C. L. Maier, and J. S. Pober, “Cytomegalovirus-infected human endothelial cells can stimulate allogeneic CD4memory t cells by releasing antigenic exosomes,” The Journal of Immunology, vol. 182, pp. 1548–1559, Jan. 2009.

58. D. W. Greening, S. K. Gopal, R. Xu, R. J. Simpson, and W. Chen, “Exosomes and their roles in immune regulation and cancer,” Seminars in Cell & Developmental Biology, vol. 40, pp. 72–81, Apr. 2015.

59. I. Huang-Doran, C.-Y. Zhang, and A. Vidal-Puig, “Extracellular vesicles: novel mediators of cell communication in metabolic disease,” Trends in Endocrinology & Metabolism, vol. 28, no. 1, pp. 3–18, 2017.

60. M. R. Anderson, F. Kashanchi, and S. Jacobson, “Exosomes in viral disease,” Neurotherapeutics, vol. 13, pp. 535–546, June 2016.

61. U. Consortium, “Uniprot: a worldwide hub of protein knowledge,” Nucleic acids research, vol. 47, no. D1, pp. D506–D515, 2019.

62. P. Samaras, T. Schmidt, M. Frejno, S. Gessulat, M. Reinecke, A. Jarzab, J. Zecha, J. Mergner, P. Giansanti, H.-C. Ehrlich, S. Aiche, J. Rank, H. Kienegger, H. Krcmar, B. Kuster, and M. Wilhelm, “ProteomicsDB: a multi-omics and multiorganism resource for life science research,” Oct. 2019.

63. L. Lautenbacher, P. Samaras, J. Muller, A. Grafberger, M. Shraideh, J. Rank, S. T. Fuchs, T. K. Schmidt, M. The, C. Dallago, H. Wittges, B. Rost, H. Krcmar, B. Kuster, and M. Wilhelm, “ProteomicsDB: toward a FAIR open-source resource for life-science research,” Nucleic Acids Research, Nov. 2021.

64. J. H. M. van Gils, D. Gogishvili, J. van Eck, R. Bouwmeester, E. van Dijk, and S. Abeln, “How sticky are our proteins? quantifying hydrophobicity of the human proteome,” Bioinformatics Advances, p. vbac002, jan 2022.

65. P. J. A. Cock, T. Antao, J. T. Chang, B. A. Chapman, C. J. Cox, A. Dalke, I. Friedberg, T. Hamelryck, F. Kauff, B. Wilczynski, and M. J. L. de Hoon, “Biopython: freely available python tools for computational molecular biology and bioinformatics,” Bioinformatics, vol. 25, pp. 1422–1423, Mar. 2009.

66. J. Lobry and C. Gautier, “Hydrophobicity, expressivity and aromaticity are the major trends of amino-acid usage in 999 escherichia coli chromosome-encoded genes,” Nucleic acids research, vol. 22, no. 15, pp. 3174–3180, 1994.

67. K. Guruprasad, B. B. Reddy, and M. W. Pandit, “Correlation between stability of a protein and its dipeptide composition: a novel approach for predicting in vivo stability of a protein from its primary sequence,” Protein Engineering, Design and Selection, vol. 4, no. 2, pp. 155–161, 1990.

68. J. Kyte and R. F. Doolittle, “A simple method for displaying the hydropathic character of a protein,” Journal of molecular biology, vol. 157, no. 1, pp. 105–132, 1982.

69. B. K. Bhandari, P. P. Gardner, and C. S. Lim, “Solubility-weighted index: fast and accurate prediction of protein solubility,” Bioinformatics, vol. 36, no. 18, pp. 4691–4698, 2020.

70. N. de Groot, I. Pallarés, F. X. Avilés, J. Vendrell, and S. Ventura BMC Structural Biology, vol. 5, no. 1, p. 18, 2005.

71. C. J. Sigrist, L. Cerutti, N. Hulo, A. Gattiker, L. Falquet, M. Pagni, A. Bairoch, and P. Bucher, “Prosite: a documented database using patterns and profiles as motif descriptors,” Briefings in bioinformatics, vol. 3, no. 3, pp. 265–274, 2002.

72. N. Hulo, A. Bairoch, V. Bulliard, L. Cerutti, B. A. Cuche, E. De Castro, C. Lachaize, P. S. Langendijk-Genevaux, and C. J. Sigrist, “The 20 years of prosite,” Nucleic acids research, vol. 36, no. suppl 1, pp. D245–D249, 2007.

73. C. J. Sigrist, E. De Castro, L. Cerutti, B. A. Cuche, N. Hulo, A. Bridge, L. Bougueleret, and I. Xenarios, “New and continuing developments at prosite,” Nucleic acids research, vol. 41, no. D1, pp. D344–D347, 2012.

74. E. De Castro, C. J. Sigrist, A. Gattiker, V. Bulliard, P. S. Langendijk-Genevaux, E. Gasteiger, A. Bairoch, and N. Hulo, “Scanprosite: detection of prosite signature matches and prorule-associated functional and structural residues in proteins,” Nucleic acids research, vol. 34, no. suppl 2, pp. W362–W365, 2006.

75. D. Wang, Y. Liang, and D. Xu, “Capsule network for protein post-translational modification site prediction,” Bioinformatics, vol. 35, pp. 2386–2394, Dec. 2018.

76. D. Wang, D. Liu, J. Yuchi, F. He, Y. Jiang, S. Cai, J. Li, and D. Xu, “MusiteDeep: a deep-learning based webserver for protein post-translational modification site prediction and visualization,” Nucleic Acids Research, vol. 48, pp. W140–W146, Apr. 2020.

77. A. Krogh, B. Larsson, G. Von Heijne, and E. L. Sonnhammer, “Predicting transmembrane protein topology with a hidden markov model: application to complete genomes,” Journal of molecular biology, vol. 305, no. 3, pp. 567–580, 2001.

78. P. V. Hornbeck, B. Zhang, B. Murray, J. M. Kornhauser, V. Latham, and E. Skrzypek, “PhosphoSitePlus, 2014: mutations, PTMs and recalibrations,” Nucleic Acids Research, vol. 43, pp. D512–D520, Dec. 2014.

79. H. Huang, C. N. Arighi, K. E. Ross, J. Ren, G. Li, S.-C. Chen, Q. Wang, J. Cowart, K. Vijay-Shanker, and C. H. Wu, “iPTMnet: an integrated resource for protein post-translational modification network discovery,” Nucleic Acids Research, vol. 46, pp. D542–D550, Nov. 2017.

80. M. Blanc, F. David, L. Abrami, D. Migliozzi, F. Armand, J. Bürgi, and F. G. van der Goot, “SwissPalm: Protein palmitoylation database,” F1000Research, vol. 4, p. 261, July 2015.

81. M. Blanc, F. P. A. David, and F. G. van der Goot, “SwissPalm 2: Protein s-palmitoylation database,” in Methods in Molecular Biology, pp. 203–214, Springer New York, 2019.

82. F. Pedregosa, G. Varoquaux, A. Gramfort, V. Michel, B. Thirion, O. Grisel, M. Blondel, P. Prettenhofer, R. Weiss, V. Dubourg, J. Vanderplas, A. Passos, D. Cournapeau, M. Brucher, M. Perrot, and Édouard Duchesnay, “Scikit-learn: Machine learning in python,” Journal of Machine Learning Research, vol. 12, no. 85, pp. 2825–2830, 2011.

83. S. M. Lundberg and S.-I. Lee, “A unified approach to interpreting model predictions,” Advances in neural information processing systems, vol. 30, 2017.

84. B. Dhondt, E. Geeurickx, J. Tulkens, J. Van Deun, G. Vergauwen, L. Lippens, I. Miinalainen, P. Rappu, J. Heino, P. Ost, et al., “Unravelling the proteomic landscape of extracellular vesicles in prostate cancer by density-based fractionation of urine,” Journal of extracellular vesicles, vol. 9, no. 1, p. 1736935, 2020.

85. J. A. Martínez-Greene, K. Hernández-Ortega, R. Quiroz-Baez, O. Resendis-Antonio, I. Pichardo-Casas, D. A. Sinclair, B. Budnik, A. Hidalgo-Miranda, E. Uribe-Querol, M. del Pilar Ramos-Godínez, and E. Martínez-Martínez, “Quantitative proteomic analysis of extracellular vesicle subgroups isolated by an optimized method combining polymer-based precipitation and size exclusion chromatography,” Journal of Extracellular Vesicles, vol. 10, apr 2021.

86. Z. Xie, A. Bailey, M. V. Kuleshov, D. J. Clarke, J. E. Evangelista, S. L. Jenkins, A. Lachmann, M. L. Wojciechowicz, E. Kropiwnicki, K. M. Jagodnik, et al., “Gene set knowledge discovery with enrichr,” Current protocols, vol. 1, no. 3, p. e90, 2021.

