## Supplemental Material for "Proteome encoded determinants of protein sorting into extracellular vesicles"

#### Supporting Figures

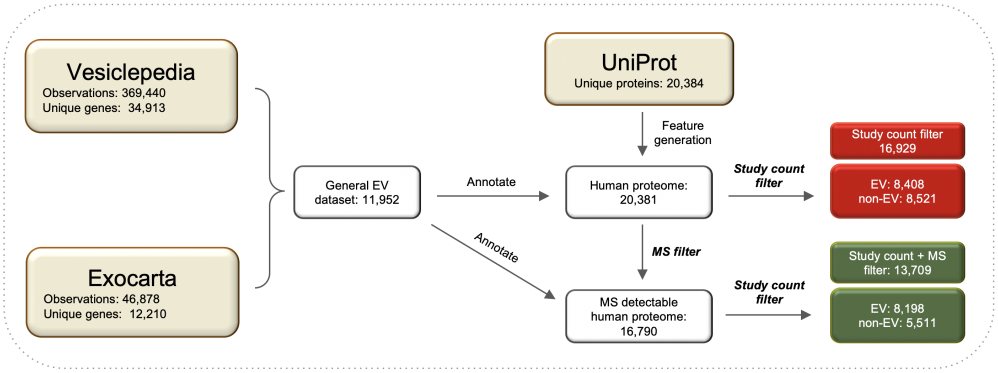


**Figure S1. Data curation workflow for less stringently filtered datasets.** Two alternative datasets were generated by a similar data curation workflow as described for the discovery set albeit some filtering steps were skipped. The general EV dataset was used to annotate the human proteome to obtain the study count filtered set (red). The MS-detectable human proteome was annotated by the general EV dataset leading to the study count and MS-filtered set (green). Both datasets are less stringently filtered than the discovery set (Figure 1) as no isolation workflow filtering was performed. EV - extracellular vesicle, MS - mass spectrometry.)

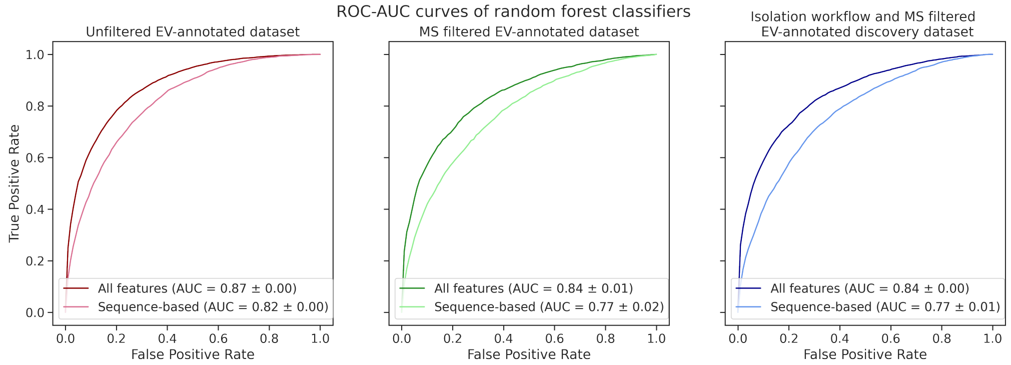


**Figure S2. Performance of random forest models trained on less stringently filtered datasets.** Random forest models trained on datasets that skipped some of the filtering steps show similar performance for EV association prediction. The unfiltered dataset (red) performs slightly better indicating that this classifier effectively (also) predicts MS detectability. Including the isolation workflow filter does not affect the model performance significantly.

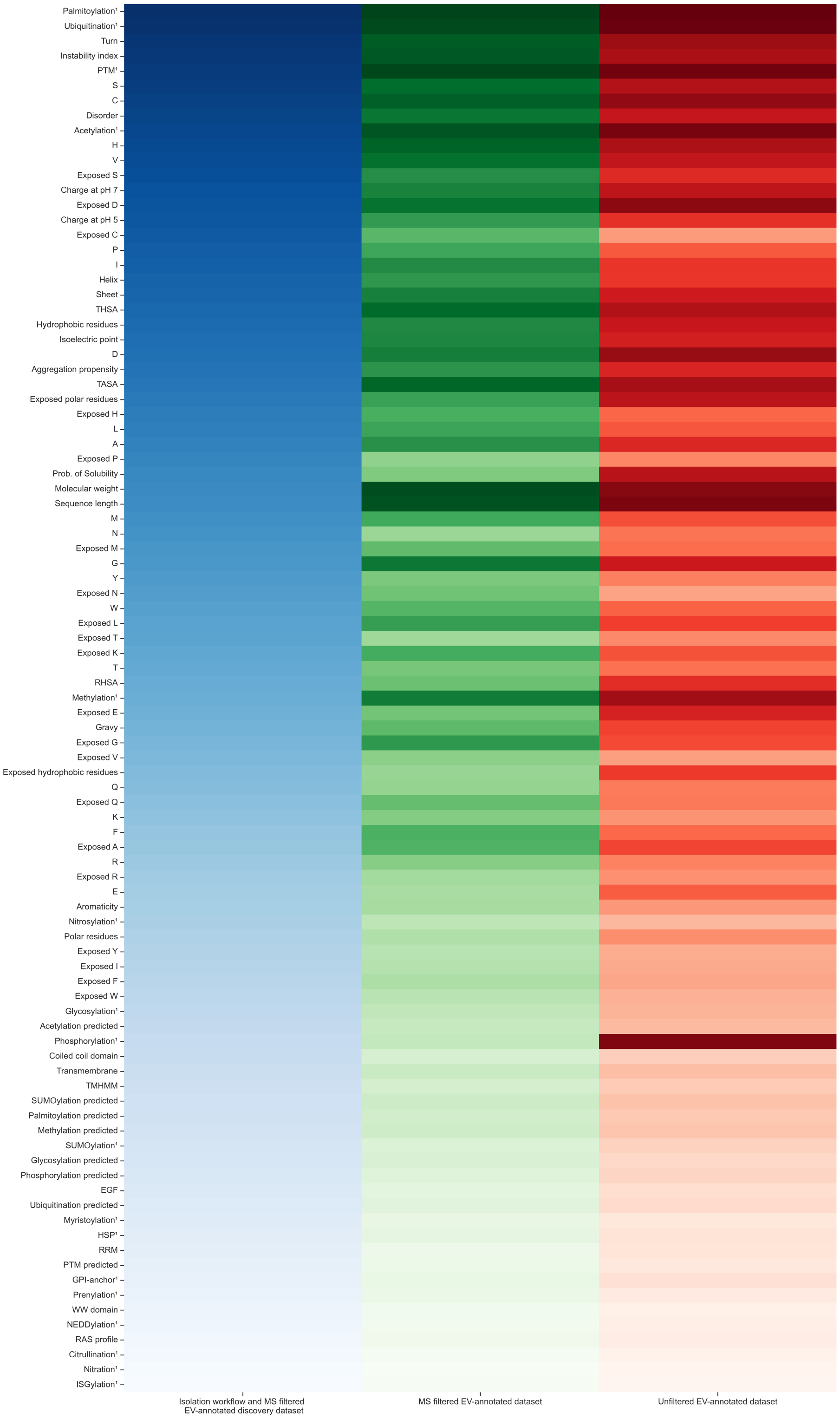


**Figure S3. Heat map of feature importance ranking.** The feature importances of the three RF models are ranked. The RFs have been trained on datasets of differing stringency, i.e., unfiltered (red), MS-filtered (green) and isolation and MS-filtered datasets. Marked features ($^1$) are curated annotations. MS - mass spectrometry, RF - random forest.

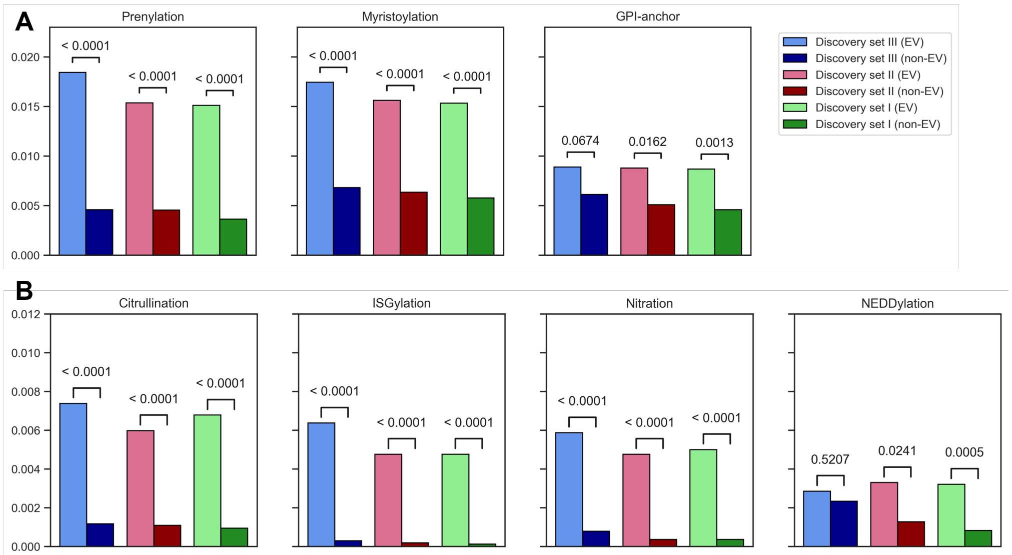


**Figure S4. Uncommon PTM annotations across the EV and non-EV proteome.** The fraction of annotated proteins regarding less common PTM types show stark differences between EV and non-EV classes. All included PTM types are enriched in the EV class but the absolute number of annotations is scarce. EV - extracellular vesicle, PTM - post-translational modification.

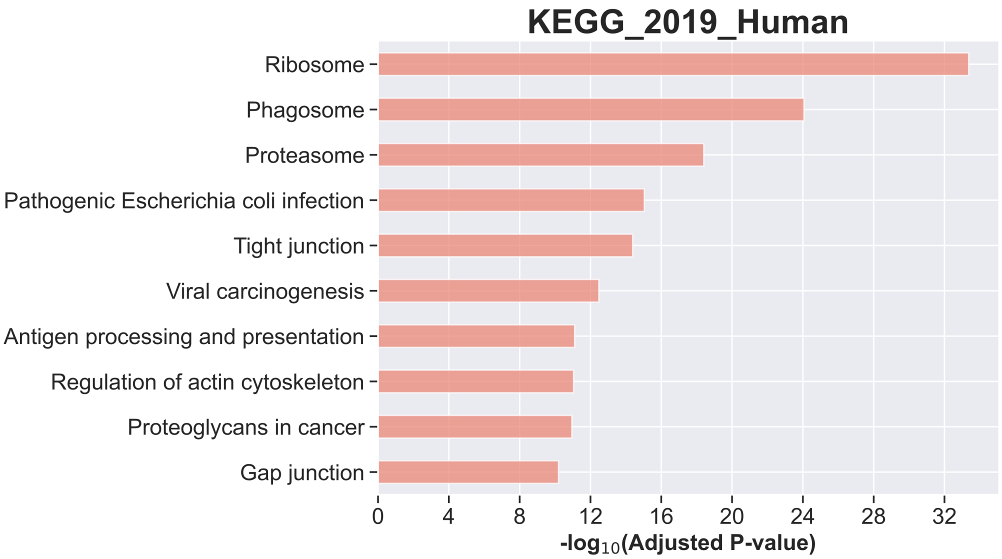


**Figure S5. Enrichment analysis of the most common EV proteins.** Proteins that are often identified in EVs (top EV proteins with occurrences in at least 30 different studies) were selected from our discovery set III and analysed with GSEApy enrichr tool (KEGG 2019 Human library) [1]. KEGG pathways that are enriched include ribosome, phagosome, proteasome, and pathways associated with infection and immune response.

### Supporting Tables

**Table S1.** Description of all generated sequence-based features implemented in the random forest classifiers.

| Feature | Source | Comments |
|---|---|---|
| Sequence length | Protein sequence | Sum of amino acids residues |
| Molecular weight | | |
| Amino acids count | | Number of residues for 20 standard amino acids in the sequence; normalized by length |
| Polar amino acids count | | Sum of polar amino acids residues (D, E, H, K, N, Q, R, T); normalized by length |
| Hydrophobic amino acids count | | Sum of hydrophobic amino acids residues (A, C, F, L, I, M, V, W, Y); normalized by length |
| TASA | NetSurfP-2.0 [2] | Derived as a global feature, sum of ASA of all AA residues |
| THSA | | Derived as a global feature, sum of ASA of hydrophobic AA residues |
| RHSA | [3] | Derived as a global feature, RHSA = THSA / TASA |
| Secondary structure | | Derived as a global feature; residue can be designated as $\alpha$ helix, $\beta$ sheet or turn |
| Disorder | | Derived as a global feature, disorder threshold per residue = 0.4 |
| Exposed AA count | | Number of exposed residues for 20 standard AAs in the sequence; normalized by total number of exposed AAs |
| Aromaticity | Biopython module [4] | Relative frequency of F, W and Y residues |
| Instability index | [5] | An index higher than 40 indicates the protein is probably unstable |
| Gravy | [6] | Calculates the hydropathic character of a protein |
| Isoelectric point | | |
| Charge at pH 7 | | |
| Charge at pH 5 | | |
| Probability of solubility | SoDoPe [7] | Scoring of a protein's solubility on a scale of 0 to 1; soluble proteins score higher |
| Aggregation propensity | [8] | Aggregation propensities were derived for each AA based on experimental data |
| Coiled coil domain | PROSITE [9] | Motif identifiers: PS51842; PS51376; PS51338; PS50067; PS50027; PS00029; PS51649; PS51121; PS51370; PS51776; PS51386; PS51649; PS51121; PS51370; PS51776; PS51386; PS51131; PS51663; PS00914; PS50151; PS50119 |
| EGF | | Motif identifier: PS50026; PS00022; PS01186 |
| RAS profile | | Motif identifier: PS51421 |
| RRM | | Motif identifier: PS50102 |
| WW domain | | Motif identifiers: PS51035; PS01179; PS50020 |
| PTM (prediction) | MusiteDeep | Prediction of any PTM |
| Acetylation (prediction) | [10] | Predictions of N6-acetyllysine |
| Glycosylation (prediction) | | Predictions of N-linked glycosylation, O-linked glycosylation |
| Methylation (prediction) | | Methylarginine, Methyllysine |
| Palmitoylation (prediction) | | Predictions of S-palmitoylation-cysteine |
| Phosphorylation (prediction) | | Predictions of Phosphoserine/threonine, Phosphotyrosine |
| SUMOylation (prediction) | | Prediction of SUMOylation |
| Ubiquitination (prediction) | | Prediction of Ubiquitination |

**Table S2.** Description of all annotation features implemented in the random forest classifiers.

| Feature | Database | Comments |
|---|---|---|
| PTM | UniProt [11] | Keyword: PTM |
| Acetylation | UniProt<br>PhosphositePlus [12]<br>iPTM [13] | Keyword: Acetylation |
| Glycosylation | PhosphoSitePlus<br>iPTM | Combined annotations of O-GlcNAc and O-GalNAc<br>Combined annotations of N-Glycosylation, O-Glycosylation, C-Glycosylation, and S-Glycosylation |
| Methylation | UniProt<br>PhosphositePlus<br>iPTM | Keyword: Methylation |
| Palmitoylation | UniProt<br>SwissPalm [14] | Keyword: Palmitate |
| Phosphorylation | UniProt<br>PhosphoSitePlus<br>iPTM | Keyword: Phosphoprotein |
| SUMOylation | PhosphoSitePlus<br>iPTM | |
| Ubiquitination | PhosphoSitePlus<br>iPTM | |
| Nitrosylation | UniProt<br>iPTM | Keyword: S-nitrosylation |
| GPI-anchor | UniProt | Keyword: GPI-anchor |
| Myristoylation | UniProt<br>iPTM | Keyword: Myristate |
| Prenylation | UniProt | Keyword: Prenylation |
| Citrullination | UniProt | Keyword: Citrullination |
| Nitration | UniProt | Keyword: Nitration |
| NEDDylation | UniProt | Text mining of PTM commentary |
| ISGylation | UniProt | Text mining of PTM commentary |
| Transmembrane protein | UniProt | Keyword: Transmembrane |
| Heat shock protein | UniProt | Keyword: Heat shock protein |

11. U. Consortium, "Uniprot: a worldwide hub of protein knowledge," *Nucleic acids research*, vol. 47, no. D1, pp. D506–D515, 2019.

12. P. V. Hornbeck, J. M. Kornhauser, S. Tkachev, B. Zhang, E. Skrzypek, B. Murray, V. Latham, and M. Sullivan, "PhosphoSitePlus: a comprehensive resource for investigating the structure and function of experimentally determined post-translational modifications in man and mouse," *Nucleic Acids Research*, vol. 40, pp. D261–D270, Dec. 2011.

13. H. Huang, C. N. Arighi, K. E. Ross, J. Ren, G. Li, S.-C. Chen, Q. Wang, J. Cowart, K. Vijay-Shanker, and C. H. Wu, "iPTMnet: an integrated resource for protein post-translational modification network discovery," *Nucleic Acids Research*, vol. 46, pp. D542–D550, Nov. 2017.

14. M. Blanc, F. P. A. David, and F. G. van der Goot, "SwissPalm 2: Protein s-palmitoylation database," in *Methods in Molecular Biology*, pp. 203–214, Springer New York, 2019.
